# Distinct protein synthesis requirements for coupled excitatory and inhibitory long-term co-plasticity in mouse hippocampus

**DOI:** 10.64898/2026.09.15.751693

**Authors:** Jadwiga Jabłońska, Dominika Orzoł, Dominika Ciurko, Grzegorz Wiera, Jerzy W. Mozrzymas

**Author notes:** **Correspondence:** (G.W.), (J.W.M). These authors contributed equally (Jadwiga Jabłońska, Dominika Orzoł). **Author contributions:** Conceptualization: J.J., G.W., and J.W.M.; Investigation and data analysis: J. J., D. O., D. C., and G. W.; Supervision: G.W. and J.W.M.; Funding acquisition: J.W.M.; Writing – Original Draft Preparation: J. J., D. O., G. W., J. W. M.; Writing – Review & Editing: J.J., D.O., G.W., J.W.M. **Declaration of interests:** The authors declare no competing interests.

## Abstract

The functioning of neuronal networks critically depends on the coordinated interaction between excitation and inhibition. The properties of glutamatergic and GABAergic synapses are finely tuned by network activity, and the heterosynaptic nature of inhibitory plasticity further emphasizes their functional coupling. Although the dependence of glutamatergic plasticity on protein synthesis is well established, the role of translation in inhibitory plasticity remains unclear. Herein, we investigated how protein synthesis supports the concurrent plasticity at inhibitory and excitatory inputs onto CA1 pyramidal neurons. To this end, we applied a high-frequency stimulation protocol of CA3-CA1 input combined with prolonged postsynaptic depolarization of principal cells. This induced both glutamatergic long-term potentiation (LTP) and inhibitory LTP (iLTP) at synapses formed by somatostatin-expressing interneurons onto pyramidal neurons. Although LTP was observed in slices from both juvenile (≤45 days) and adult (>45 days) mice, iLTP was only present in the adult group, revealing an unusual developmental profile. Moreover, in adult animals, excitatory LTP was required for the induction of iLTP but did not determine its magnitude. We found that bath application of translation inhibitor cycloheximide (CHX) strongly attenuated LTP and converted co-occurring iLTP into iLTD. However, when CHX was restricted to the recorded postsynaptic neuron, the initial phase of iLTP proceeded normally, but GABAergic currents gradually returned to baseline levels, indicating intact induction but impaired maintenance. In contrast, postsynaptic CHX left excitatory LTP unaffected. Together, these findings identify two distinct translational mechanisms: a postsynaptic one necessary for strengthening inhibition and a non-postsynaptic process required to potentiate excitation.

## Introduction

GABAergic inhibition in the brain exerts sophisticated control over neural network activity at several levels. It shapes brain rhythms that underlie various aspects of cognition. At the neuronal level, it controls the gain, spike timing, and dendritic integration of principal excitatory neurons and can also target other interneurons, causing disinhibition (Kullander and Topolnik, 2021). The balance between excitation and inhibition (E/I) that underlies these functions is not static but is continuously fine-tuned by activity-dependent plasticity at both synapse classes (Vogels et al., 2011; Field et al., 2020; Cupolillo et al., 2024). These two forms of plasticity are not independent, as potentiation of excitatory inputs can drive concurrent changes at neighboring inhibitory synapses, heterosynaptically coupling inhibitory adjustment to excitatory long-term change (Hennequin et al., 2017; Ravasenga et al., 2022; Agnes and Vogels, 2024). The induction requirements of this co-dependent GABAergic plasticity (iLTP/iLTD) are increasingly well characterized (Wu et al., 2022; Wiera et al., 2024; Jabłońska et al., 2025; Lech et al., 2026), but the mechanisms that maintain it remain far less understood than those at glutamatergic synapses. In particular, the maintenance phase of excitatory plasticity depends critically on *de novo* protein synthesis (Frey et al., 1988; Sutton and Schuman, 2006; Di Prisco et al., 2014), but whether this process sustains co-dependent inhibitory plasticity onto pyramidal cells is unknown. Additionally, the polarity of GABAergic signaling shifts from depolarizing to hyperpolarizing during early postnatal life (Ben-Ari et al., 2007), and glutamatergic plasticity follows its own age-dependent trajectory showing greater efficacy at earlier developmental stages (Monfort and Felipo, 2007; Le et al., 2022). Whether long-term GABAergic plasticity itself follows a comparable developmental course has remained largely unexplored.

Protein synthesis is indeed required for inhibitory plasticity, but it is distributed across several cellular compartments. In dissociated cultures, iLTP induced by bath NMDA requires dendritic synthesis of GABAA receptor subunits and of the scaffolding protein gephyrin. This process is regulated by activity-dependent de-repression of microRNAs (miR-376c and miR-153) downstream of the calcineurin–NFAT– HDAC pathway (Rajgor et al., 2020; Welle et al., 2024). Presynaptically expressed endocannabinoid-mediated iLTD, by contrast, requires protein synthesis in the axonal bouton of the CB1-expressing interneuron, rather than in the postsynaptic cell (Younts et al., 2016), where it drives structural remodeling of the terminal (Monday et al., 2020). Beyond GABAergic plasticity, protein synthesis in astrocytes tunes the threshold for excitatory LTP and shifts the E/I balance, in part by weakening inhibitory transmission onto pyramidal cells (Sharma et al., 2023). In addition to its role in activity-dependent plasticity, translational machinery sets the baseline strength of inhibition. The eukaryotic elongation factor 2 (eEF2) kinase, which arrests translation elongation, acts as a brake on GABAergic transmission, and its inhibition upregulates presynaptic synapsin-2b and postsynaptic α5-GABAA receptors and strengthens inhibition (Heise et al., 2017; Beretta et al., 2022). Likewise, cap-dependent translation initiated via the eIF4E/4E-BP2 pathway controls the expression of the inhibitory adhesion molecule neuroligin-2, thereby influencing the E/I balance (Gkogkas et al., 2013). Finally, mTORC1 dysregulation restricted to interneurons reduces synaptic inhibition of pyramidal cells (Haji et al., 2020). Protein synthesis therefore governs inhibitory transmission at multiple loci and through distinct mechanisms.

What remains unaddressed is how protein synthesis supports the co-occurrence of plasticity at the inhibitory and excitatory inputs onto pyramidal neurons. Evidence for translation-dependent inhibitory plasticity comes almost entirely from chemical induction protocols, in which bath-applied NMDA induces inhibitory changes in dissociated neurons, while excitatory synaptic responses were not monitored (Rajgor et al., 2020; Welle et al., 2024). Such protocols therefore cannot address how the two forms of plasticity are related and, in particular, how they depend on protein synthesis. Here, we characterized an age-dependent, heterosynaptic form of iLTP at inhibitory synapses from somatostatin-expressing interneurons onto pyramidal cells (SST→PC), induced by LTP of excitatory CA3→CA1 input. The SST→PC projection was chosen because GABAergic synapses are formed on PC dendrites, a compartment that expresses postsynaptic inhibitory plasticity (Wiera and Mozrzymas, 2026) and supports local translation (Biever et al., 2019). It is also noteworthy that GABAergic synapses onto SST-positive interneurons show prominent NMDA-induced iLTP (Brzdąk et al., 2023, 2025) and NMDA-dependent upregulation of tonic inhibition (Wyroślak et al., 2023). Additionally, learning increases protein synthesis, particularly in this interneuron subpopulation, alongside similar changes in PCs, with a mechanism dependent on eIF2α (Sharma et al., 2020). Thus, SST-positive interneurons express GABAergic plasticity, whether they target pyramidal neurons, are innervated by GABAergic cells, or participate in tonic inhibition. We found that a bath-applied translation inhibitor markedly attenuated LTP and converted co-occurring iLTP to iLTD, whereas the same inhibitor restricted to the recorded postsynaptic neuron abolished iLTP while leaving LTP intact. These results reveal two protein synthesis mechanisms that operate in parallel: a postsynaptic one necessary for potentiation of inhibition, and a non-postsynaptic process required for potentiation of excitation. This partitioning implies that blocking protein synthesis does not simply weaken plasticity but biases its outcome toward excitation, a consideration that applies to any experiment in which translation is globally inhibited.

## Results

### Co-expression of iLTP in SST→PC synapses is age-dependent

Before addressing the role of protein synthesis in GABAergic and excitatory plasticity, we characterized both phenomena across a wide range of ages and in both sexes, as these factors are known to shape synaptic plasticity (Le et al., 2022). Previous studies on GABAergic heterosynaptic plasticity, including our own, have relied on transient NMDA application (Marsden et al., 2007; Petrini et al., 2014; Jabłońska et al., 2024; Lech et al., 2026). Here, we employed a more physiological approach, in which prolonged postsynaptic depolarization was paired with the stimulation of glutamatergic synaptic input. This strategy is not only closer to the physiological conditions of plasticity induction but also evokes inhibitory and excitatory plasticity in the same neuron with the same protocol, allowing us to directly characterize their co-occurrence.

Before characterizing synaptic plasticity, we verified the specificity of the SST-Cre driver line used throughout this study. Immunostaining for RFP in an SST-Cre:Ai14 reporter cross confirmed reporter expression restricted to somatostatin-positive interneurons in stratum oriens (Supplemental Fig. 1.)

Excitatory and inhibitory currents (EPSCs and IPSCs) were simultaneously recorded from the same CA1 pyramidal cells in hippocampal slices. EPSCs were evoked by stimulation of the CA3→CA1 pathway and measured at −70 mV, close to the reversal potential of the inhibitory current. Simultaneously, IPSCs were optogenetically evoked (light stimulation of SST INs) and recorded at −60 mV. To induce excitatory LTP, the cell was held at 0 mV while CA3→CA1 afferents were stimulated at 100 Hz for 1 s, and this paradigm was repeated three times at 20 s intervals (3×100Hz @0mV). This protocol produced robust excitatory LTP and induced concurrent potentiation of SST → PC inhibition in the same cells (iLTP, Fig. 1A,D). This approach allowed us to follow the two forms of plasticity as they were induced simultaneously in the same neuron.

**Figure 1.**
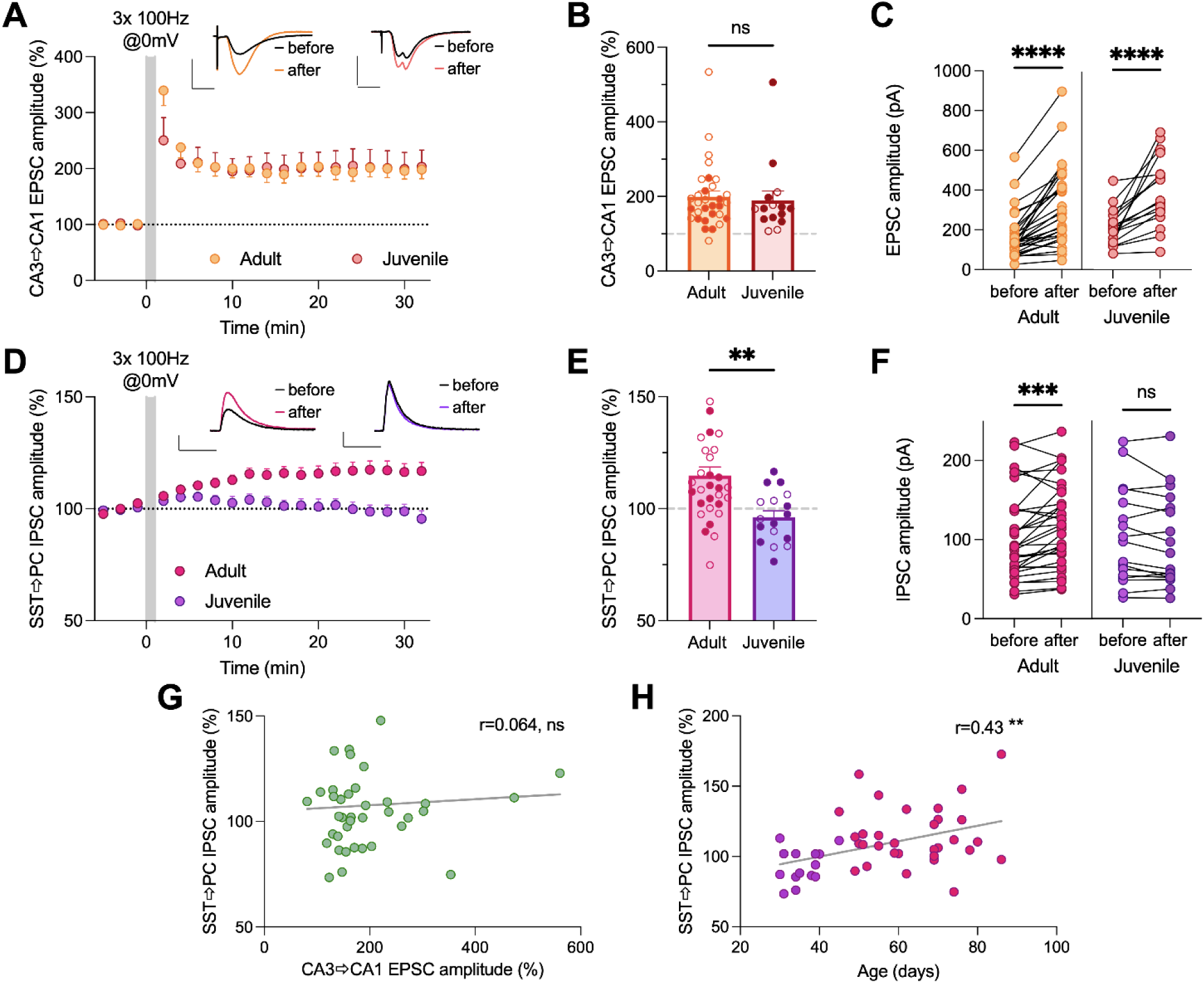
Induction of CA3→CA1 excitatory LTP leads to age-dependent concurrent plastic changes at SST→PC synapses. **(A)** Time course of normalized CA3→CA1 PC EPSC amplitudes recorded in brain slices isolated from juvenile (P≤45) and adult (P>45) mice following the induction of synaptic plasticity with a 3×100 Hz @0mV stimulation (gray shading). Representative averaged EPSC traces recorded before (black) and 25–30 min (colored) after induction protocol are shown. **(B)** Summary of normalized EPSC amplitudes measured 25–30 min after 100 Hz stimulation in both age groups. Juvenile, *n* = 17 cells; adult, *n* = 31 cells; unpaired *t-test*. **(C)** Paired plots showing EPSC amplitudes measured before stimulation and 25–30 min later for individual recordings in each experimental group (Wilcoxon’s test). **(D)** Time course of normalized SST→PC IPSC amplitudes recorded in the same cells as in (A). Representative averaged IPSC traces recorded before (black) and 25–30 min after (colored) plasticity induction are shown. **(E)** Summary of normalized IPSC amplitudes measured 25–30 min after plasticity induction. Juvenile, *n* = 17 cells; adult, *n* = 31 cells; unpaired *t-test*. **(F)** Paired plots showing IPSC amplitudes measured before stimulation and 25–30 min later for individual recordings in adult and juvenile groups (Wilcoxon test). **(G)** Correlation analysis between the magnitude of CA3→CA1 LTP and concurrent SST→PC plasticity expressed in the same CA1 PCs for all recordings. No significant correlation was detected (Pearson’s r = 0.064, p = 0.70). **(H)** Correlation between animal age and the magnitude of SST→PC iLTP. The magnitude of iLTP increased with age, indicating enhanced inhibitory synaptic plasticity in older animals (Pearson’s r = 0.43, P = 0.0034). **** P < 0.0001; ** P < 0.01; ns – nonsignificant. Filled points in **B** and **E** indicate data from female mice

Excitatory LTP was consistently present across all ages examined (Fig. 1 B), whereas iLTP was variable and absent in a substantial fraction of cells (Fig. 1E). Grouping animals as juveniles (P≤45) and adults (P>45) revealed a clear difference in the extent of iLTP (adult: 108.4 ± 18.4 pA before, 121.8 ± 16.8 pA after, p = 0.0009; juvenile: 132.6 ± 17.3 pA before, 126.1 ± 17.7 pA after, p = 0.17; Wilcoxon tests; Fig. 1D-E). Thus, in the adult group, LTP induction was reliably accompanied by SST→PC iLTP, whereas in the juvenile group, no lasting inhibitory change occurred (juvenile: 93.4 ± 3.3 % of baseline; adult: 114.8 ± 3.8 %; p = 0.0017; unpaired *t*-test; Fig. 1E). This age dependence was confirmed at the level of individual recordings, where the magnitude of iLTP increased with age (Pearson’s r = 0.43, p = 0.0034; Fig. 1G). The P45 threshold defining adult mice was set a priori, consistent with our previous studies using this convention (Wiera and Mozrzymas, 2026). Excitatory plasticity, by contrast, was similar in both groups (juveniles: 196.3 ± 18.0 %; adults: 188.3 ± 26.4 %; p = 0.73; *t*-test; Fig. 1B-C), indicating that the age difference was specific to inhibition. Although iLTP required excitatory LTP, its magnitude did not scale with the magnitude of LTP (Pearson’s r = 0.064, p = 0.70; Fig. 1H), indicating that excitatory potentiation gates the inhibitory change rather than setting its size. We found no differences between the sexes (Supplemental Figs. 2 and 3). Because iLTP emerged reliably in adult animals, mice older than P45 were used in all subsequent experiments; this developmental characterization served primarily to define the experimental model in which co-expressed excitatory and inhibitory LTP could be reliably studied.

### Bath cycloheximide reduces excitatory LTP and converts coupled iLTP to iLTD

To test whether protein synthesis is required for excitatory LTP and SST→PC iLTP co-expressed in the same CA1 pyramidal cell, we first applied cycloheximide (CHX) to the bath through the recording, blocking translation throughout the slice. In interleaved vehicle recordings, the 3×100Hz protocol potentiated both inhibitory (IPSC: 107 ± 23.6 pA before, 121.7 ± 28.7 pA after, p = 0.04; Fig. 2A,B) and excitatory inputs (EPSC: 115.4 ± 16.3 pA before, 365.9 ± 61.2 pA after, p = 0.005; Fig. 2D,E). With CHX in the bath, excitatory LTP was still present, although its extent was markedly smaller than that under control conditions (132.1 ± 19.6 pA before, 188.6 ± 34.5 pA after, p = 0.02; Fig. 2D,E). When normalized to the baseline, excitatory LTP decreased from 342.2 ± 62.3 % in vehicle to 144.5 ± 12.8 % under CHX (p = 0.003, Fig. 2F). In the presence of CHX, SST→PC IPSCs were not potentiated by the 3×100Hz protocol but instead underwent iLTD (107.4 ± 16.9 pA before, 89.6 ± 11.1 pA after, p = 0.047; Fig. 2A,B). Normalized inhibitory plasticity fell from 112.3 ± 2.9 % in vehicle to 89.8 ± 6.3 % under CHX (p = 0.01, Fig. 2C), confirming a switch from iLTP to iLTD. These data show that both excitatory and inhibitory forms of co-expressed plasticity depend on de novo protein synthesis, but the extent of this dependence shows qualitative and quantitative differences.

**Figure 2.**
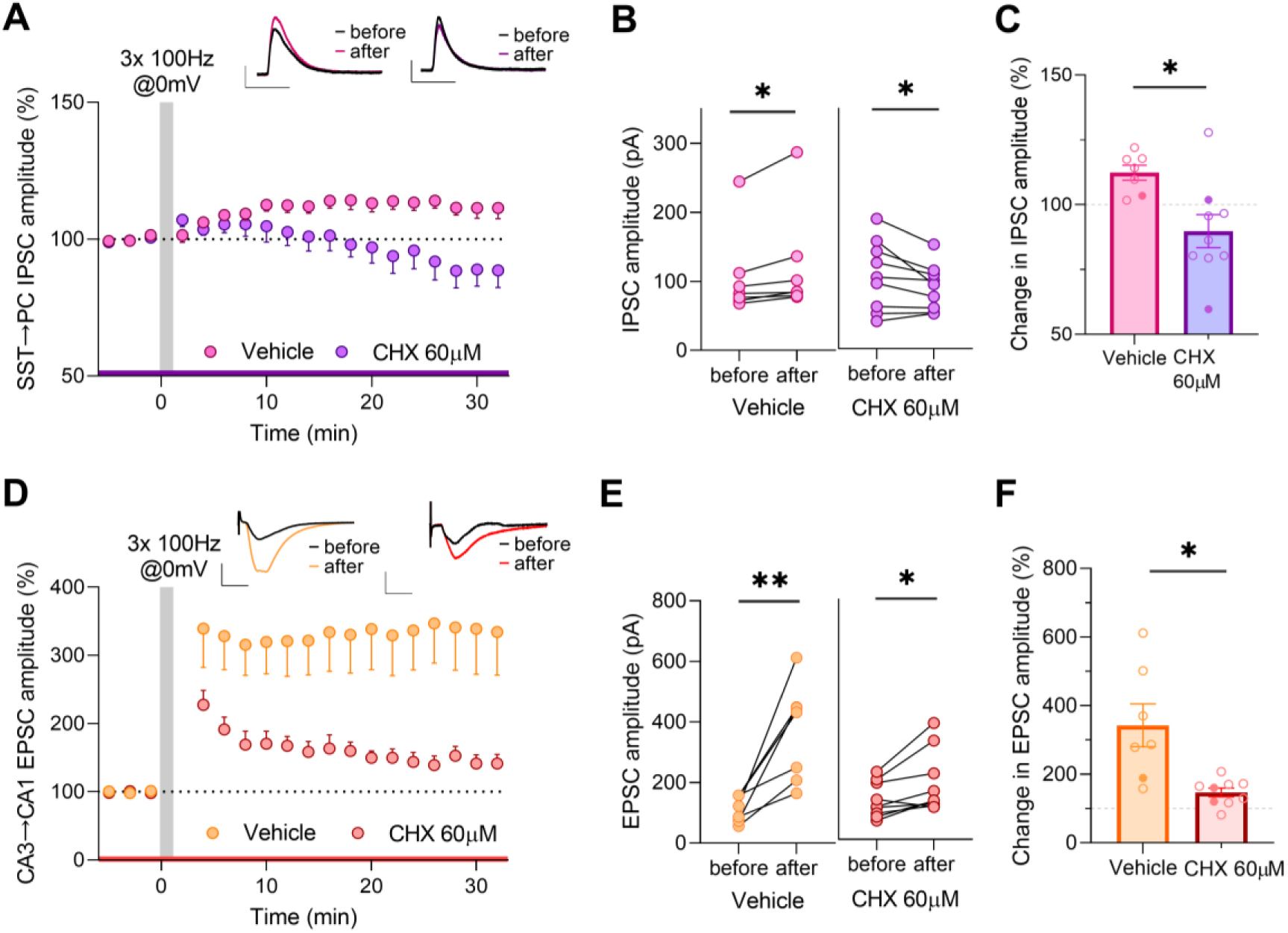
Inhibition of protein synthesis attenuates CA3→CA1 excitatory LTP and converts SST→PC iLTP to iLTD. **(A)** Time course of normalized SST→PC IPSC amplitudes before and after the induction of synaptic plasticity with a 3 × 100 Hz stimulation protocol (gray shading). Representative averaged IPSC traces recorded before (black) and 25–30 min after (colored) plasticity induction are shown above for the vehicle and cycloheximide (CHX) groups. Scale bars: 50 ms, 50 pA. The horizontal line in the color of the corresponding trace indicates the period of vehicle or CHX presence (60 µM). **(B)** Paired plots showing IPSC amplitudes measured before induction and 25–30 min after the plasticity protocol for individual cells recorded in vehicle and CHX groups. **(C)** Summary of normalized IPSC amplitudes measured after plasticity induction, demonstrating a switch from SST→PC iLTP to iLTD in the presence of CHX. **(D)** Time course of normalized CA3→CA1 EPSC amplitudes recorded concurrently from the same CA1 PCs, as in (A). Representative averaged EPSC traces recorded before (black) and after (colored) plasticity induction. Scale bars: 10 ms, 100 pA. **(E)** Paired plots showing EPSC amplitudes measured before and after the plasticity protocol in individual cells. **(F)** Summary of normalized EPSC amplitudes after plasticity induction, showing that CHX significantly reduced the magnitude of CA3→CA1 excitatory LTP. Filled points in **C** and **F** indicate data from female mice. Data are presented as mean ± SEM. Vehicle with DMSO, n = 7 cells; CHX, n = 9 cells. Statistical analyses: **B** and **E**, paired t-test; **C** and **F**, unpaired t-test. * P < 0.05; ** P < 0.01.

### Postsynaptic cycloheximide blocks iLTP while leaving LTP intact

Bath-applied CHX showed that protein synthesis contributes to both CA3→CA1 excitatory LTP and SST→PC iLTP, although it affects them differently (Fig. 2). Bath application, however, blocks translation everywhere in the slice, in presynaptic and postsynaptic neuronal compartments, and in non-neuronal cells, such as astrocytes and microglia, any of which could contribute to plasticity. Because the potentiation of inhibition observed here is expected to be postsynaptic (Lech et al., 2026; Wiera and Mozrzymas, 2026), we limited the protein synthesis block to the recorded pyramidal cell by adding CHX only to the pipette solution. We first tested whether intracellular CHX altered the basal IPSC amplitude. Throughout the recording, intracellular CHX did not change SST→PC IPSC amplitude (vehicle: 82.0 ± 9.7 pA before, 82.2 ± 9.5 pA after, p = 0.9; CHX: 108.5 ± 20.9 pA before, 110.5 ± 14.7 pA after, p = 0.8; Fig. 3A,C).With vehicle in the pipette, the 3×100 Hz protocol potentiated SST→PC IPSCs (78.6 ± 9.8 pA before, 102.8 ± 14.3 pA after, p = 0.0006; Fig. 3B). In contrast, SST → PC iLTP was abolished when CHX was added to the pipette (107.4 ± 13.1 pA before, 108.0 ± 12.6 pA after, p = 0.91; Figs. 3B–D). However, the inhibitory current amplitude still rose transiently over the first 10 min after plasticity induction (112 ± 3.4 % at 10 min, p = 0.003 vs. baseline, paired *t*-test; Fig. 3B) but eventually decayed to the baseline. This indicates that the induction of inhibitory plasticity proceeded normally over the first 10 min minutes or so but could not be maintained without postsynaptic protein synthesis.

**Figure 3.**
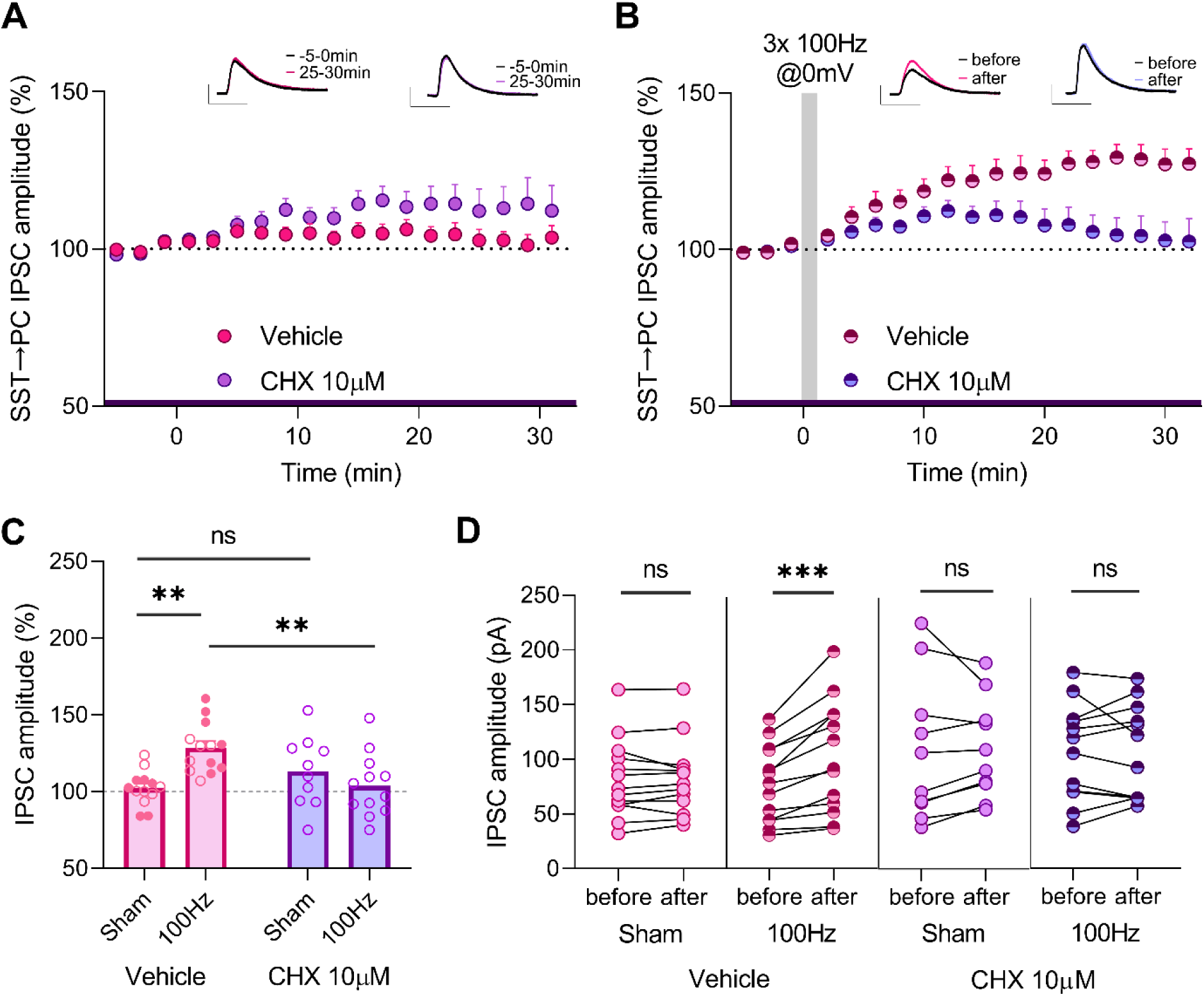
Intracellular inhibition of protein synthesis in the PC abolishes SST→PC inhibitory long-term potentiation. **(A)** Baseline recordings (sham without 3×100Hz protocol) of the time course of normalized SST→PC IPSC amplitudes recorded with vehicle pipette solution (vehicle) and with cycloheximide (CHX) that was present throughout the recording only in the pipette solution. Representative averaged IPSC traces from the first 5 min (black) and from 25–30 min (colored) of recording are shown above. The horizontal line in the color of the corresponding trace indicates the period of vehicle or CHX presence (10 µM). **(B)** Time course of normalized SST→PC IPSC amplitudes following the induction of synaptic plasticity with a 3×100Hz stimulation protocol (gray shading) in recordings obtained with vehicle intracellular solution (vehicle) or with intracellular CHX. Representative averaged IPSC traces recorded before (black) and 25–30 min after (colored) plasticity induction are shown. **(C)** Summary of normalized IPSC amplitudes measured 25–30 min after sham or 3 × 100 Hz stimulation under control conditions (vehicle) and with intracellular CHX, demonstrating that intracellular CHX prevents SST→PC iLTP. Vehicle-Sham, *n* = 13 cells; Vehicle-100 Hz, *n* = 13 cells; CHX-Sham, *n* = 10 cells; CHX-100Hz, *n* = 12 cells. Two-way ANOVA (stimulation × pipette solution): stimulation, F(1, 44) = 2.71, p = 0.10 (4.4% of total variance); pipette solution, F(1, 44) = 1.7, p = 0.20 (2.8%); interaction, F(1, 44) = 11.7, p = 0.0014 (19% of total variance), followed by Tukey’s post hoc multiple-comparison test. Filled points indicate data from female mice. **(D)** Paired plots showing IPSC amplitudes measured during the first 5 min and 25–30 min later for individual recordings in each experimental group (paired *t-test*). Scale bar (representative traces): 50ms, 50pA. *** P < 0.001; ** P < 0.01; ns – nonsignificant.

### Postsynaptic cycloheximide spares excitatory LTP that drives iLTP

We next asked whether postsynaptic CHX, which abolished iLTP (Fig. 3), also affected co-expressed excitatory LTP induced by the same protocol in the same CA1 pyramidal cells. With CHX present in the pipette solution, the basal EPSC amplitude drifted slightly upward (232.9 ± 39.8 pA before, 297.3 ± 55.8 pA after, p = 0.02), and this drift did not differ from recordings with the vehicle solution in the pipette (112.2 ± 9.1 % vs. 122.5 ± 9.1 %, p = 0.99, two-way ANOVA with Tukey’s post hoc correction for multiple comparisons; Fig. 4A). Therefore, we did not interpret this as a genuine change in basal excitatory transmission. With the vehicle in the pipette, the 3×100 Hz protocol potentiated the excitatory pathway alongside iLTP (EPSC: 194.6 ± 36.5 pA before, 376.1 ± 56.5 pA after, p = 0.0001; Fig. 4B–D). Unexpectedly, with CHX in the pipette, excitatory LTP was still induced (203.3 ± 37.7 pA before, 397.7 ± 52.4 pA after; p = 0.001; Fig. 4D). Its magnitude did not differ significantly from that with vehicle solution (248.1 ± 41.5 % with CHX vs. 217.5 ± 29.8 % with vehicle, p = 0.85, two-way ANOVA with Tukey post hoc test; Fig. 4B–D), even though iLTP in the same cells was abolished (Fig. 3B). These experiments demonstrated that confining the protein synthesis block to the recorded neuron separated the two plasticities, preventing potentiation of inhibition while leaving potentiation of excitation intact.

**Figure 4.**
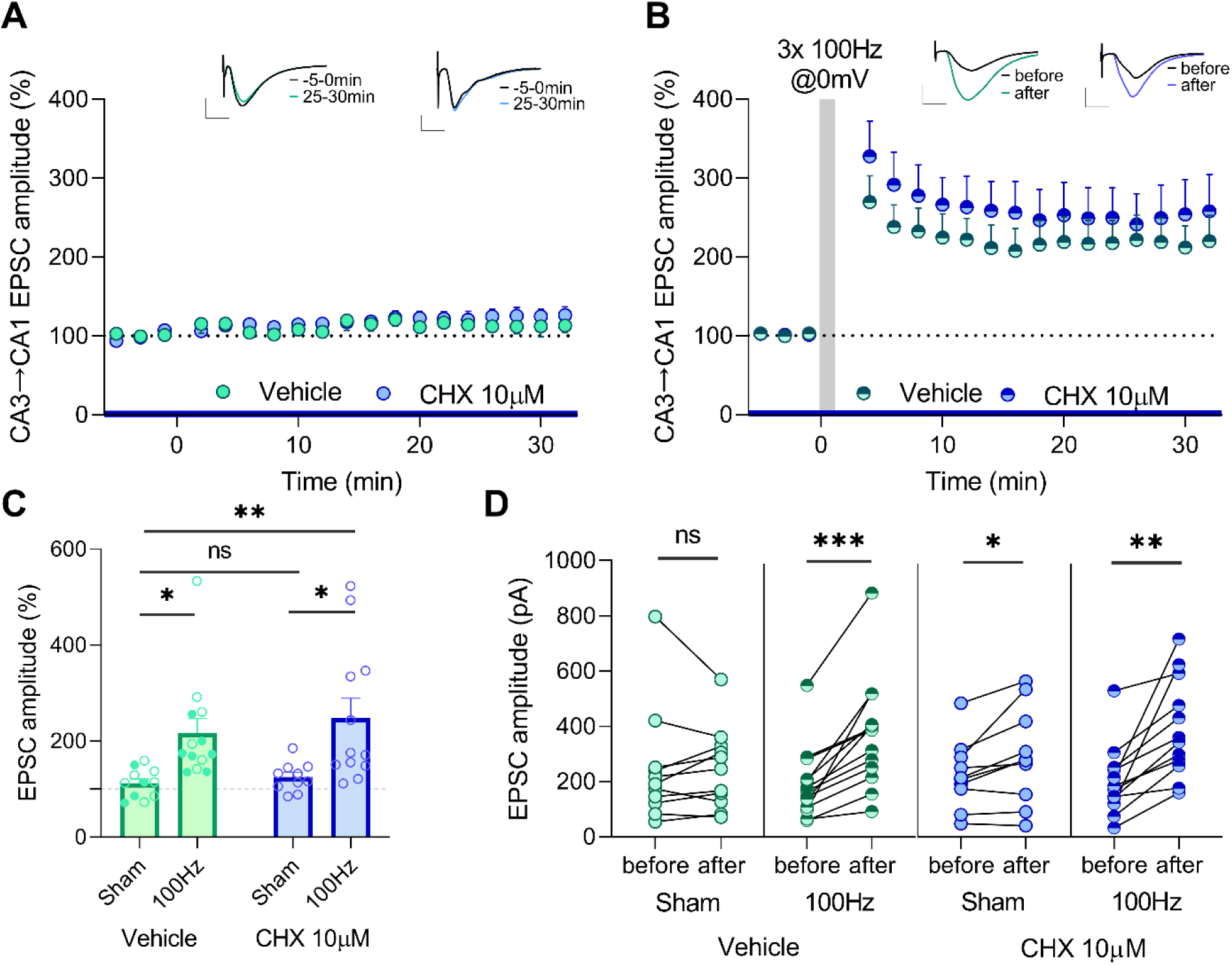
Intracellular inhibition of protein synthesis does not affect CA3→CA1 excitatory long-term potentiation. **(A)** Baseline recordings of the time course of normalized CA3→CA1 EPSC amplitudes obtained with a pipette solution containing vehicle or cycloheximide (CHX). Sample sizes: Vehicle, n = 11 cells; CHX, n = 10 cells. Representative averaged EPSC traces from the first 5 min (black) and from 25–30 min (colored) of the recording are shown above. The horizontal line in the color of the corresponding trace indicates the period of vehicle or CHX presence (10 µM). **(B)** Time course of normalized CA3→CA1 EPSC amplitudes following the induction of synaptic plasticity with a 3 × 100 Hz stimulation protocol (gray shading) in recordings obtained with the control intracellular solution or intracellular CHX. Sample sizes: Vehicle, n = 13 cells; CHX, n = 12 cells. Representative averaged EPSC traces recorded before (black) and 25–30 min after (colored) plasticity induction are shown. **(C)** Summary of normalized EPSC amplitudes measured 25–30 min after sham or 100 Hz stimulation in control and intracellular CHX conditions, demonstrating that intracellular CHX does not alter the magnitude of CA3→CA1 eLTP. Two-way ANOVA (stimulation × pipette solution): stimulation, F(1, 42) = 16.3, p = 0.0002 (27.6 % of total variance); pipette solution, F(1, 42) = 0.59, p = 0.47 (1.0 %); interaction, F(1, 42) = 0.098, p = 0.76 (0.16 % of total variance), followed by Tukey’s post hoc multiple-comparison test. Filled points indicate data from female mice. **(D)** Paired plots showing EPSC amplitudes measured before stimulation and 25–30 min later for individual recordings in each experimental group (paired *t* tests). Data are presented as mean ± SEM. *P* < 0.05; *P* < 0.01; *P* < 0.001; ns, not significant. Scale bars (representative traces): 10 ms, 200 pA. *** P < 0.001; ** P < 0.01; * P < 0.05; ns – nonsignificant.

## Discussion

The major finding of the present study is that co-occurring excitatory and inhibitory long-term plasticity, elicited using the same protocol and recorded from single CA1 pyramidal cells, differ in their sensitivity to protein synthesis blockade. When CHX was applied to the bath, iLTP was converted to iLTD, while LTP was reduced but not abolished. When the blocker was restricted to the postsynaptic pyramidal neuron, the late phase of iLTP was eliminated without conversion to iLTD, whereas LTP was spared. The co-expression of excitatory and inhibitory synaptic plasticity described here is consistent with previous studies (Ravasenga et al., 2022; Lech et al., 2026; Wiera and Mozrzymas, 2026). Importantly, SST input is highly amenable to GABAergic plasticity, more than e.g. PV synapses (Sharma et al., 2020; Jabłońska et al., 2026; Wiera and Mozrzymas, 2026). Moreover, SST-INs are known to increase protein synthesis both *in vivo* after learning (Sharma et al., 2020) and *in vitro* after chemical and TBS stimulation (Honoré et al., 2022); therefore, their plasticity is expected to be particularly prone to factors affecting protein synthesis. Thus, potentiation of inhibition depends on protein synthesis both within the postsynaptic cell and outside it, whereas excitatory LTP depends on synthesis beyond the recorded pyramidal cell. These findings demonstrate that although, in the considered model, inhibitory and excitatory plasticities are induced by the same protocol in the same cell, their dependence on protein synthesis differs, most likely reflecting distinct molecular pathways. Notably, a subset of cells expressed robust LTP without accompanying iLTP, indicating that excitatory potentiation is necessary but insufficient for inhibitory changes.

The finding that CHX in the postsynaptic neuron does not affect excitatory LTP elicited by our 3 × 100Hz protocol coupled with prolonged postsynaptic depolarization may appear surprising, as it is commonly accepted that long-lasting synaptic plasticity and related memory processes are associated with protein synthesis (Frey et al., 1988; Klann and Sweatt, 2008; Costa-Mattioli et al., 2009). Several studies have shown, however, that the late phase of excitatory potentiation can be expressed without new proteins (e.g., Abbas et al., 2009; Villers et al., 2012). Similar rules apply to GABAergic neuroplasticity. For instance, Field et al. (2021) have shown that driving GABAA receptors into the desensitized conformation, together with their phosphorylation by PKC, is sufficient to produce iLTP. These observations reflect the principle that synaptic plasticity (both excitatory and inhibitory), depending on various induction paradigms and physiological conditions, relies on a wide spectrum of distinct mechanisms, and protein synthesis is just one of them, and its impact may depend on other players. Such a broader view finds direct support in the work of Fonseca et al., (2006), who showed that late LTP was preserved when protein synthesis and proteasome-dependent degradation were blocked together, indicating that L-LTP depends on the combined action of synthesis and degradation of plasticity proteins rather than on synthesis alone.

### Postsynaptic protein synthesis maintains heterosynaptic iLTP

Intracellular CHX left basal SST→PC transmission intact, yet abolished iLTP, indicating a key role of protein synthesis in the induced plastic changes rather than in supporting IPSCs under basal conditions. Moreover, the inhibitory amplitude continued to rise over the first few minutes after the induction protocol and then decayed to baseline, indicating intact induction and a selective failure of maintenance. This assigns the new protein synthesis requirement of iLTP to a consolidation step downstream of induction, in agreement with the biphasic scenario described for chemically induced NMDA-iLTP, where early receptor trafficking is translation-independent and later maintenance requires new proteins (Rajgor et al., 2020; Welle et al., 2024). This is in line with our previous observations that SST→PC iLTP, when co-expressed with excitatory LTP, undergoes a consolidation phase that requires the reorganization of neuroligin-2-dependent synaptic adhesion (Lech et al., 2026) and extracellular proteolysis (Wiera et al., 2024). Inhibitory potentiation, like excitatory LTP, therefore resolves into an induction phase and a translation-dependent consolidation phase. Our recordings address the early maintenance of this consolidation over the first ∼30 min after induction; whether postsynaptic synthesis is also required to sustain iLTP over longer periods remains to be determined.

Prior evidence that inhibitory potentiation requires protein synthesis came from chemically induced plasticity, in which bath NMDA drove GABAergic change in dissociated neurons, but no accompanying excitatory plasticity was recorded (Rajgor et al., 2020; Welle et al., 2024). These studies demonstrated that iLTP maintenance involves dendritic synthesis of GABAA receptor subunits and gephyrin, which requires downregulation of miR-376c and miR-153 downstream of the calcineurin–NFAT–HDAC pathway. Our findings confirm that protein synthesis is crucial for iLTP and extend this requirement to the physiological induction protocol in brain slices. It remains to be determined whether calcium sources other than NMDARs, such as voltage-gated calcium channels (VGCCs), contribute to this process. The strong postsynaptic depolarization used for induction, combined with high-frequency stimulation of glutamatergic input, offers a plausible entry point for the underlying signaling pathway. During depolarization, NMDA receptor activation and possibly VGCC opening may gate a calcium signal and the ensuing dendritic translation through dephosphorylation of eEF2, a switch that couples synaptic activity to local protein synthesis (Sutton et al., 2007), and controls GABAergic transmission (Heise et al., 2017; Beretta et al., 2022). Whether this mechanism or the miRNA-dependent program accounts for the protein synthesis requirement observed here remains to be tested, but both predict a synthesis step confined to postsynaptic pyramidal neurons.

### Excitatory LTP requires protein synthesis outside the postsynaptic neuron

Compared to iLTP, excitatory LTP shows a different sensitivity profile to protein synthesis blockade. It remained unchanged when translation was blocked in the recorded pyramidal neuron and was reduced only when CHX was added to the bath. The protein synthesis supporting full expression of this LTP therefore lies outside the postsynaptic cell. Astrocytic translation is a strong candidate for this mechanism: eIF2α-dependent synthesis in astrocytes sets the threshold for hippocampal LTP and shifts the E/I balance (Sharma et al., 2023), and would be accessible to bath but not to intracellular CHX. Several other possibilities can be hypothesized. Blocking translation throughout the slice would also affect microglia and their metabolic coupling with neurons (Adler et al., 2026). Inhibiting protein synthesis in the entire slice could also reduce the synthesis and release of BDNF, which regulates several forms of synaptic plasticity (De Vincenti et al., 2019; Mei et al., 2011).

The molecular mechanism by which protein synthesis blockade throughout the slice attenuates LTP in our model remains unknown. Local axonal translation, linked to presynaptic NMDA receptors and mTOR, sustains high-frequency transmission at excitatory synapses within minutes, and bath application of a translation inhibitor acutely depresses burst transmission through a presynaptic mechanism (Wong et al., 2024). Because our LTP is induced by repeated 100 Hz stimulation, part of the plasticity reduction in the presence of bath CHX could reflect a weakened high-frequency drive during induction, rather than a postsynaptic block of the maintenance phase. However, the time course of this effect argues against this hypothesis. Potentiation in the presence of bath CHX was normal immediately after tetanization and declined thereafter, a pattern that fits a consolidation deficit rather than acute suppression of the induction drive. A comparison with dopamine-gated LTP can be instructive. This form requires postsynaptic synthesis and is blocked by intracellular translation inhibitors, whereas conventional LTP is spared (Fuchsberger et al., 2022, 2025). Our excitatory LTP behaves like conventional LTP with respect to the postsynaptic compartment, yet retains a non-postsynaptic translational dependence.

### Cycloheximide impact on plasticity balance

The two routes of CHX delivery affected inhibitory and excitatory co-plasticity to different degrees. With the block confined to the pyramidal neuron, iLTP was reduced to baseline, and eLTP was intact. With translation blocked throughout the slice, the inhibitory response fell below the baseline, whereas excitatory potentiation was weakened but preserved. In both conditions, the balance shifted toward excitation because translation blockade reduced inhibitory plasticity more severely than excitatory plasticity. What drives the depression seen under bath CHX remains unresolved; a presynaptic, endocannabinoid-dependent mechanism is unlikely, since it would require axonal synthesis that bath CHX blocks (Younts et al., 2016; Monday et al., 2020).

We admit as a limitation that sex distribution was not balanced across all experimental groups, precluding formal statistical testing for sex differences. Where sample sizes permitted (vehicle groups, Figs. 3–4), no significant sex difference in plasticity magnitude was detected. Our conclusions regarding the non-postsynaptic protein-synthesis dependence of excitatory LTP are therefore established primarily for males, and confirmation in females remains an important direction for future work.

### Developmental emergence of inhibitory potentiation

An intriguing discovery of our study is that inhibitory potentiation was present in adults but absent in juveniles. This implies that the developmental emergence of the observed inhibitory plasticity is reversed with respect to major neuroplasticity phenomena, including sensitive periods (Hensch and Fagiolini, 2004; Knudsen, 2004), neuroregeneration (Bradke, 2022), and several forms of glutamatergic LTP (Monfort and Felipo, 2007; Le et al., 2022). The major form of GABAergic plasticity, the switch from depolarizing to hyperpolarizing GABA, also occurs predominantly during early development (Ben-Ari et al., 2007). Thus, our observations extend our knowledge of the extraordinary versatility of GABAergic plasticity in the developmental context. Notably, the excitatory LTP measured here was similar across the entire age range examined, which also differs from the previously described greater efficacy in earlier developmental stages (Monfort and Felipo, 2007; Le et al., 2022). Thus, it is noteworthy that the “reversed” developmental gate that acts on inhibitory plasticity is ineffective on the excitatory drive, whose plasticity remains constant despite the same induction protocol. Several maturational processes could set this developmental profile for the iLTP described here, including the refinement of SST→PC connectivity, release of growth and modulatory factors, changes in chloride handling that alter the driving force and sign of GABAergic signaling, and maturation of the dendritic translation program itself. Translational regulators that constrain synapse number and composition are developmentally downregulated over this period, and the miRNA and mTOR pathways controlling local translation are themselves under developmental control (Randolph et al., 2024). We emphasize that a developmental change in translational capacity is only one of several plausible explanations considered above for the age-dependence of iLTP, and it is not implied by our finding that postsynaptic protein synthesis is required for iLTP in adults (Fig. 3); these remain two separate observations that the present study does not mechanistically link.

### Behavioral implications and conclusions

Systemic inhibition of protein synthesis during learning produces a striking dissociation. It blocks natural memory recall, but optogenetic activation of engram cells still retrieves the memory, defining a “silent engram” (Ryan et al., 2015; Roy et al., 2017). Under bath CHX, excitatory potentiation was reduced but preserved in direction, whereas inhibitory potentiation was lost and reversed toward depression. This asymmetry raises the possibility that the global translation blockade does not act uniformly across a circuit but affects GABAergic and glutamatergic synapses differently. If consolidation normally strengthens the dendritic inhibition that governs how selectively an engram is reactivated, then losing that inhibition would leave the engram accessible to direct stimulation but not to natural cues, which depend on precise input patterns.

In summary, our results show that inhibitory and excitatory plasticity, despite being evoked by the same high-frequency protocol, show different dependencies on protein synthesis, with the former being more effectively regulated by this process. This implies that the signal generated as the demand for new proteins is partitioned across cellular compartments, allowing excitation and inhibition to be potentiated together while remaining under separate, translational control.

## Materials and Methods

### Ethical approval and animals

All animal procedures used in this study were in agreement with Polish law regulations (Act on the Protection of Animals Used for Scientific or Educational Purposes, January 15, 2015, with subsequent amendments) and EU Directive 2010/63/EU. Experimental procedures involving genetically modified organisms were approved by the Polish Ministry of the Environment (approval no. 69/2023).

Experiments used mice expressing channelrhodopsin-2 in SST-positive interneurons generated by crossing homozygous Sst-IRES-Cre (Ssttm2.1(cre)Zjh/J; RRID:IMSR_JAX: 013044) and Ai32 (B6.Cg-Gt(ROSA)26Sortm32(CAGCOP4*H134R/EYFP)Hze/J; RRID:IMSR_JAX:024109). Except for the developmental characterization (where P35-P120 mice were used, Fig. 1), all recordings were obtained from adult mice aged P45-120. Animals were housed at the Animal Facility of Wroclaw Medical University (Poland) in standard cages under a 12-hour light/dark cycle with same-sex littermates. Water and food were available *ad libitum*.

### Preparation of acute brain slices

Acute transverse slices (350 µm) were prepared as described by Ting et al. (2018) with minor modifications (Jabłońska et al., 2024; Wiera et al., 2024). Holding aCSF (in mM: 92 NaCl, 2.5 KCl, 1.2 NaH₂PO₄, 30 NaHCO₃, 20 HEPES, 25 glucose, 5 sodium ascorbate, 2 thiourea, 3 sodium pyruvate, 1.3 MgSO₄, 2.5 CaCl₂) and recovery NMDG solution (in mM: 93 NMDG, 2.5 KCl, 1.2 NaH₂PO₄, 30 NaHCO₃, 20 HEPES, 25 glucose, 5 sodium ascorbate, 2 thiourea, 3 sodium pyruvate, 10 MgSO₄, 0.5 CaCl₂) were continuously bubbled with carbogen (95% O₂ / 5% CO₂). Briefly, mice were anesthetized with isoflurane (5% in oxygen), and brains were rapidly dissected, mounted with cyanoacrylate glue, and sectioned using a vibratome (VT1200, Leica) in ice-cold holding aCSF. Slices were transferred to ∼200 ml of warmed (34 °C) recovery NMDG solution, and 2 M NaCl was added stepwise (250 µl–2 ml) over an incubation period that was lengthened for older animals (20–35 min). A gradual increase in NaCl concentration improves slice viability and preserves membrane integrity (Ting et al., 2018). After recovery, slices were held at room temperature in holding aCSF and used within 8 h.

### Electrophysiological recordings

For patch-clamp recordings, slices were superfused (3–4 ml/min) with carbogen-saturated recording aCSF (in mM: 119 NaCl, 2.5 KCl, 1.3 NaH₂PO₄, 26 NaHCO₃, 1.3 MgSO₄, 2.5 CaCl₂, and 11 D-glucose) at 28 °C. The tissue was visualized using an upright microscope (AxioExaminer Z1, Zeiss) with 4× and 40× water-immersion objectives under IR-DIC optics.

Borosilicate pipettes (GB150TF-8P, Science Products) were pulled on a Sutter puller and had resistances of 3–5 MΩ when filled with intracellular solution (in mM: 135 K-gluconate, 5 NaCl, 5 Mg-ATP, 0.3 Na-GTP, 10 phosphocreatine, 10 HEPES; pH 7.3, 285 mOsm). SST-positive inputs were activated by 2 ms pulses of blue light from a fiber-coupled DG-4 optical switch (Sutter Instruments) with an HQ470/40× excitation filter delivered through a 40×, 1.0 NA objective. Blue light was delivered at 10 mW/mm² at the sample, a level adjusted to evoke stable, submaximal SST→PC IPSCs without failures. ChR2 expression was restricted to SST-positive interneurons by the Sst-IRES-Cre driver, so light-evoked IPSCs reflected the selective activation of SST→PC synapses. CA3→CA1 excitatory inputs were stimulated with a tungsten bipolar electrode (FHC) placed in the stratum radiatum ∼150 µm from the recorded cell. The stimulus intensity (200 µs pulses) was adjusted to evoke ∼40% of the maximal EPSC amplitude, below the threshold for a spike. Within each 10-s sweep, the cell was held at −70 mV for 600ms to record electrically evoked CA3→CA1 EPSCs, and then at −60 mV for 9.4 s to record optogenetically evoked SST→PC IPSCs. As in our previous studies (Jabłońska et al., 2024; Wiera et al., 2024; Lech et al., 2026; Wiera and Mozrzymas, 2026), the two responses were separated temporally and evoked using pathway-selective stimulation rather than pharmacological isolation. Electrical stimulation of the CA3→CA1 pathway activates glutamatergic transmission, whereas stimulation of ChR2-expressing SST interneurons via the Sst-IRES-Cre driver selectively evokes GABA release. EPSCs were recorded at −70 mV, close to the measured IPSCs reversal potential (−78.9 ± 1.1 mV, n = 101), thereby minimizing any inhibitory contribution while electrical driving force for cations was large (with reversal potential close to 0 mV). The membrane potentials are reported without correction for the liquid junction potential, which was ∼15 mV for the K-gluconate-based intracellular solution.

Signals were amplified (Multiclamp 700B), low-pass filtered at 4 kHz (Bessel), and digitized (Digidata 1550A; Molecular Devices). Data were acquired using Clampex 11.3 at a sampling rate of 20 kHz. CA1 pyramidal neurons were identified by soma morphology and location and by characteristic electrical properties (a prominent hyperpolarization-evoked sag and spike frequency adaptation). Access resistance was monitored throughout the experiment, and cells with a shift ≥20% were excluded. All recordings were performed in voltage clamp, except for firing-pattern characterization (current-clamp).

In a subset of experiments, CHX was applied either to the bath or through the pipette. For bath application, CHX was dissolved in DMSO and used at a final concentration of 60 µM (final DMSO concentration ≤ 0.1%), a concentration established to effectively block protein synthesis-dependent LTP in acute hippocampal slices while preserving baseline synaptic transmission (Ren et al., 2013). For intracellular application, CHX was dissolved in water to avoid DMSO in the pipette solution and used at a concentration of 10 µM (Fuchsberger et al., 2025).

### Plasticity induction and electrophysiological data processing

To evoke long-term plasticity of both inhibitory and excitatory inputs in the same neuron, CA3→CA1 afferents were stimulated at 100 Hz for 1 s while the postsynaptic cell was held in a voltage clamp at 0 mV; this was repeated three times at 20 s intervals.

All recorded electrophysiological signals were processed using the Clampfit 10.7 software. PSC amplitudes were averaged over 2 min bins and normalized to the mean baseline recorded before the plasticity protocol was applied. The magnitude of plasticity was measured 25–30 min after completing the protocol.

### Compounds and chemicals

Chemicals, suppliers and cat. numbers are listed in table

| Compound | Supplier | Cat. # |
| --- | --- | --- |
| CaCl<sub>2</sub> | Merck (Sigma-Aldrich) | C7902 |
| Cycloheximide | Merck (Sigma-Aldrich) | C1988 |
| D-glucose | Merck (Sigma-Aldrich) | G8270 |
| DMSO | Merck (Sigma-Aldrich) | D8418 |
| GTP | Merck (Sigma-Aldrich) | 10106399001 |
| HEPES | Merck (Sigma-Aldrich) | H4034 |
| isoflurane (Aerrane) | Baxter Polska | 8DG9623 |
| KCl | Merck (Sigma-Aldrich) | P5405 |
| K-gluconate | Merck (Sigma-Aldrich) | P1847 |
| MgATP | Merck (Sigma-Aldrich) | A9187 |
| MgSO<sub>4</sub> | Merck (Sigma-Aldrich) | M5921 |
| NaCl | Merck (Sigma-Aldrich) | S7653 |
| NaH<sub>2</sub>PO<sub>4</sub> | Merck (Sigma-Aldrich) | 71504 |
| NaHCO<sub>3</sub> | VWR | 27778.293 |
| NMDG | Merck (Sigma-Aldrich) | A3286 |
| phosphocreatine | Merck (Sigma-Aldrich) | P7936 |
| sodium ascorbate | Merck (Sigma-Aldrich) | A7631 |
| sodium pyruvate | Merck (Sigma-Aldrich) | P5280 |
| tiourea | Merck (Sigma-Aldrich) | T7875 |

### Statistical analysis

Our methodology followed several predetermined guidelines. Sample sizes were determined a priori using G*Power software, targeting a statistical power of at least 0.80. No formal statistical test for outliers was applied; all collected data points were retained without exclusion. Normality of all datasets was assessed using the Shapiro-Wilk test, and homogeneity of variance was assessed using Levene’s test; together, these determined the choice between parametric and nonparametric comparisons throughout the study. Experimental groups were assigned randomly, although the researchers were aware of the treatment conditions during the data collection and analysis phases. The data were analyzed and plotted using GraphPad Prism (v8 and v10). Normality of all datasets was assessed using the Shapiro-Wilk test. For two-group comparisons, we used Student’s t-test (paired or unpaired) for normally distributed data and the Mann-Whitney U or Wilcoxon test otherwise, based on this normality assessment and the study design. For the correlation analyses in Fig. 1G-H, normality and linearity of the data were assessed prior to selecting Pearson’s correlation coefficient. Multiple group comparisons were performed using two-way ANOVA (factors: stimulation and pipette solution) followed by Tukey’s post hoc test. Statistical significance is marked as: ns – nonsignificant; p > 0.05; * p ≤ 0.05; ** p ≤ 0.01; *** p ≤ 0.001; **** p ≤ 0.0001. Group sizes are indicated in the figure captions. n refers to the number of cells, and recordings were obtained from at least three mice per group. Each condition was tested using interleaved control and CHX recordings. Data are presented as mean ± SEM.

## Acknowledgments

This work was supported by funding from the National Science Centre (Poland) grant OPUS 2021/43/B/NZ4/01675 to J.W.M.

## Materials availability

This study did not generate any new unique reagents.

## Data and code availability

All data reported in this paper will be shared at the Polish Medical Platform (PPM, DOI: 10.60956/89t8-qq46) and upon personal request. This study did not report the original code. All data associated with this study are presented in this paper and have not been published elsewhere.

