## supplemental files for "Distinct protein synthesis requirements for coupled excitatory and inhibitory long-term co-plasticity in mouse hippocampus"

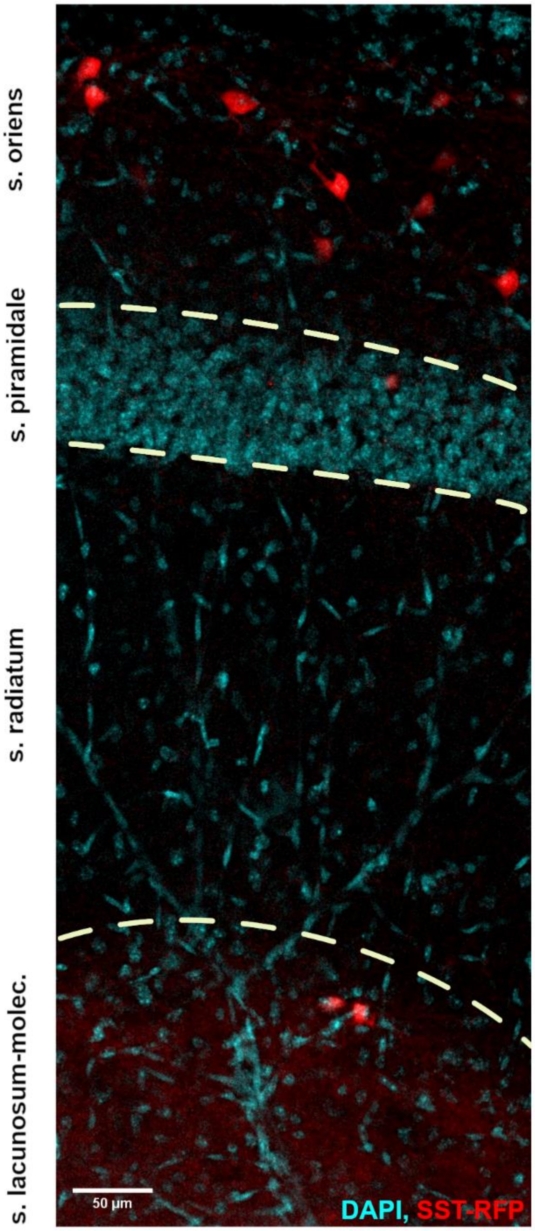


**Supplementary Figure 1. SST-Cre-driven reporter expression is restricted to somatostatin-positive interneurons in CA1.**

Representative confocal image of a transverse hippocampal section (CA1 region) from an SST-Cre:Ai14 mouse, showing DAPI (cyan) and immunodetected RFP (red, Cre-dependent tdTomato reporter). Dashed lines demarcate laminar boundaries (stratum oriens, stratum pyramidale, stratum radiatum, stratum lacunosum-moleculare). RFP-positive cell bodies are concentrated in stratum oriens, with occasional somata at the stratum radiatum/stratum lacunosum-moleculare border, consistent with the known distribution of SST-positive interneurons. Dense RFP-positive fibers are additionally observed throughout stratum lacunosum-moleculare, consistent with the characteristic axonal arborization of OLM interneurons in this layer. No RFP signal is detected within stratum pyramidale, arguing against off-target labeling of pyramidal cells. Scale bar, 50 µm.

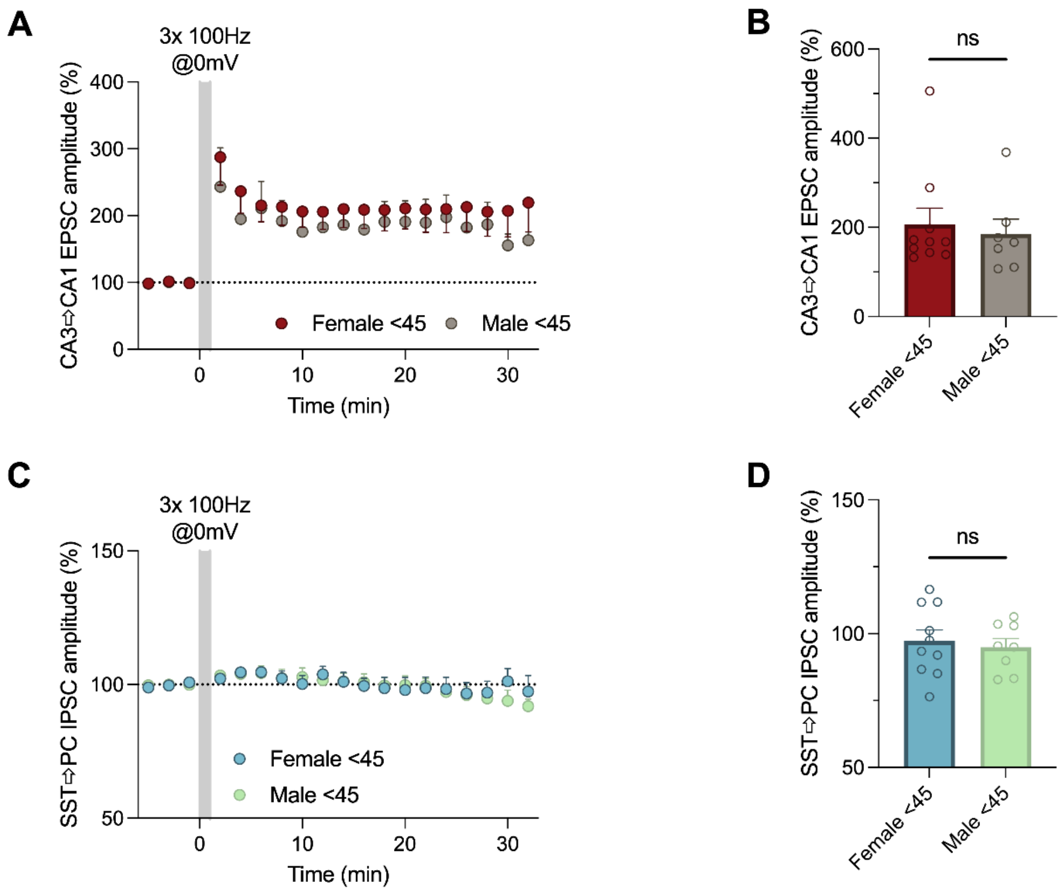


**Supplementary Figure 2. Sex does not influence CA3→CA1 excitatory synaptic plasticity or the absence of concurrent SST→PC plasticity in mice younger than P45.**

**(A)** Time course of normalized CA3→CA1 EPSC amplitudes following induction of synaptic plasticity with a 3 × 100 Hz stimulation protocol (gray shading) in female and male mice younger than P45. **(B)** Summary of normalized CA3→CA1 EPSC amplitudes measured 25–30 min after plasticity induction, showing no significant difference in LTP magnitude between female and male mice (male: 184.9 ± 21.8 %, $n$ = 7; female: 206.8 ± 51.6 %, $n$ = 10; $p$ = 0.6778, unpaired $t$ test). **(C)** Time course of normalized SST→PC IPSC amplitudes following the same 3 × 100 Hz stimulation protocol, recorded simultaneously from the same CA1 pyramidal neurons as the EPSCs shown in **A**, in female and male mice younger than P45. **(D)** Summary of normalized SST→PC IPSC amplitudes measured 25–30 min after plasticity induction, demonstrating the absence of concurrent SST→PC plasticity in both female and male mice (male: 94.9 ± 2.3 %, $n$ = 7; female: 97.2 ± 5.5 %, $n$ = 10; $p$ = 0.6770, unpaired $t$ test). Data are presented as mean ± SEM; ns, nonsignificant.

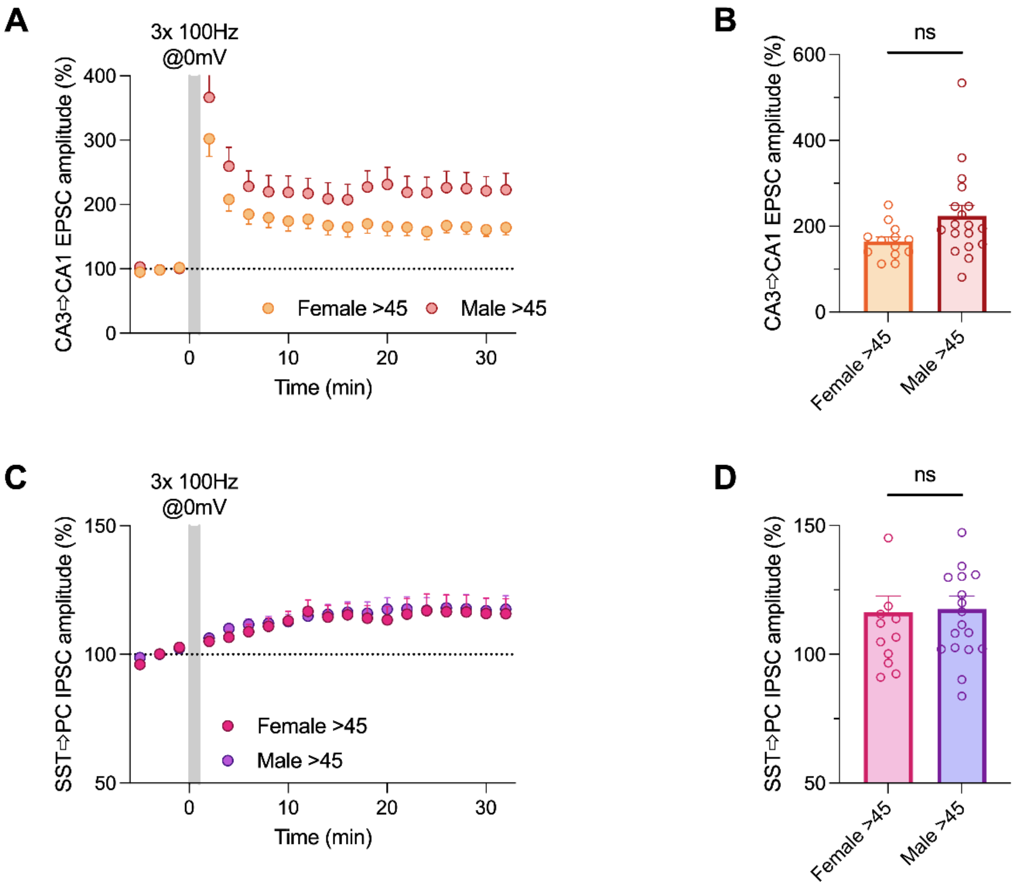


**Supplementary Figure 3. Sex does not influence CA3→CA1 excitatory synaptic plasticity or the absence of concurrent SST→PC plasticity in mice older than P45.**

**(A)** Time course of normalized CA3→CA1 EPSC amplitudes following induction of synaptic plasticity with a 3 × 100 Hz stimulation protocol (gray shading) in female and male mice older than P45. **(B)** Summary of normalized CA3→CA1 EPSC amplitudes measured 25–30 min after plasticity induction, showing no significant difference in LTP magnitude between female and male mice (male: 224.1 ± 59.7 %, $n$ = 18; female: 164.1 ± 30.1 %, $n$ = 13; $p$ = 0.0567, unpaired $t$ test). **(C)** Time course of normalized SST→PC IPSC amplitudes following the same 3 × 100 Hz stimulation protocol, recorded simultaneously from the same CA1 pyramidal neurons as the EPSCs shown in **A**, in female and male mice older than P45. **(D)** Summary of normalized SST→PC IPSC amplitudes measured 25–30 min after plasticity induction, demonstrating the induction of concurrent SST→PC iLTP in both female and male mice (male: 117.6 ± 1.3 %, $n$ = 18; female: 116.3 ± 8.1 %, $n$ = 13; $p$ = 0.8713, unpaired $t$ test). Data are presented as mean ± SEM; ns, nonsignificant.
